# Using Watershed Characteristics for Improving Fecal Source Identification

**DOI:** 10.1101/2022.06.16.496426

**Authors:** John J. Hart, Megan N. Jamison, James N. McNair, Sean A. Woznicki, Ben Jordan, Richard Rediske

## Abstract

Fecal pollution is one of the most prevalent forms of pollution affecting waterbodies worldwide, threatening public health, and negatively impacting aquatic environments. Microbial source tracking (MST) applies polymerase chain reaction (PCR) technology to help identify the source of fecal pollution. In this study, we combine spatial data for two watersheds with general and host-specific MST markers to target human, bovine, and general ruminant sources. Two different PCR technologies were applied for quantifying the targets: quantitative PCR (qPCR) and droplet digital PCR (ddPCR). We found that ddPCR had a higher detection rate (75%) of quantifiable samples compared to qPCR (27%), indicating that ddPCR is more sensitive than qPCR. The three host-specific markers were detected at all sites (n=25), suggesting that humans, cows, and ruminants are contributing to fecal contamination in these watersheds. MST results, combined with watershed characteristics, suggest that streams draining areas with low-infiltration soil groups, high septic system prevalence, and high agricultural land use are at an increased risk for fecal contamination. Microbial source tracking has been applied in numerous studies to aid in identifying the sources of fecal contamination, however these studies usually lack information on the involvement of watershed characteristics. Our study combined watershed characteristics with MST results, applying more sensitive PCR techniques, in addition to watershed characteristics to provide more comprehensive insight into the factors that influence fecal contamination in order to implement the most effective best management practices.

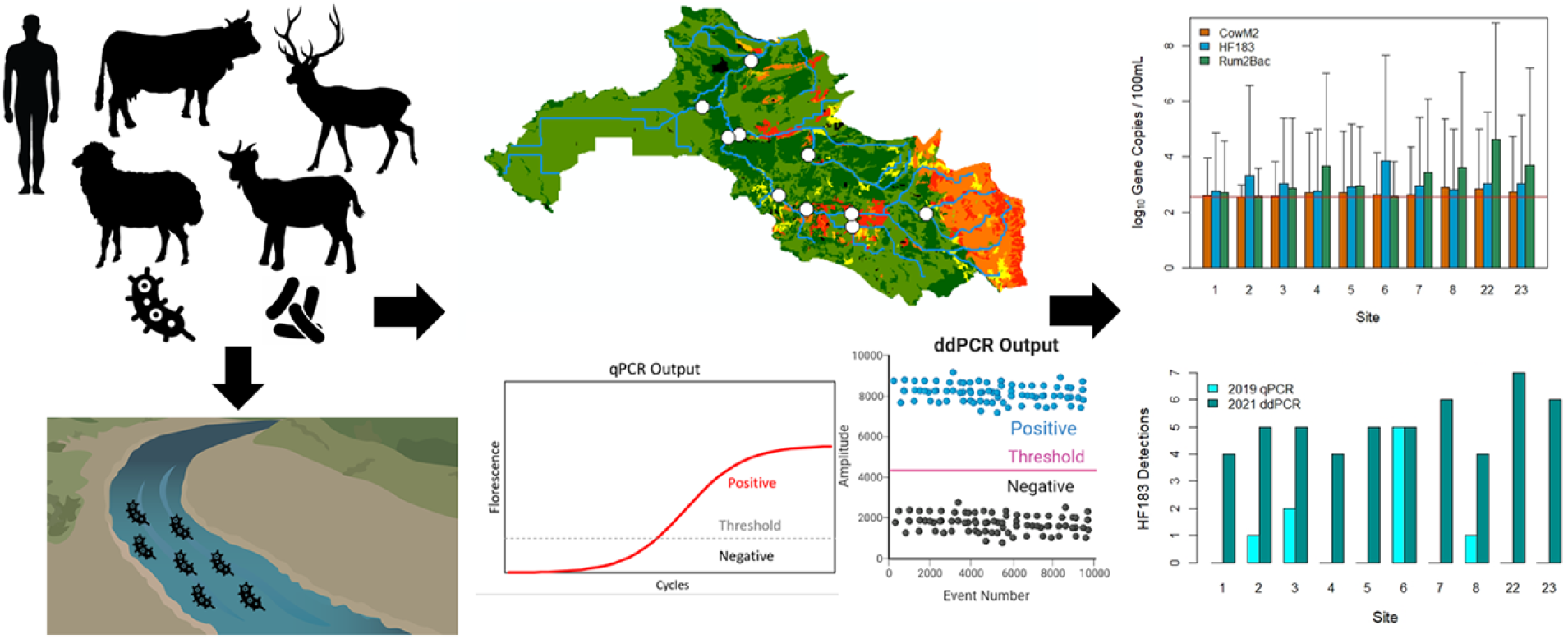

**Highlights:**

- ddPCR provided higher sensitivity over qPCR when analyzing environmental samples
- Human markers had an association with the number of septic systems in a watershed
- Every site had positive detections for all FIB markers
- Both ruminant markers were associated with low infiltration hydrologic soil groups
- Combining watershed characteristics with MST testing improved source identification

## 1. Introduction

Water quality degradation is a global problem, which is further affected by global change (Hofstra et al., 2019). The United States has over 43,000 water bodies impaired by various pollutants (US EPA, 2017). Total maximum daily load (TMDL) assessments have been conducted for many pollutants to determine the maximum amount of a pollutant allowed to enter a waterbody while remaining within the established water quality standards. The second most prevalent source of impairment is pathogen contamination from f ecal pollution, accounting for 20% of all established TMDLs (US EPA, 2017). According to the United Nations, approximately, 1.8 billion people use a water source contaminated with fecal pollution (Bain et al., 2014). Fecal pollution introduces excess nutrients and potentially harmful pathogens that pose significant risks to stream and public health (Ballesté et al., 2020; Jent et al., 2013). Ingestion of harmful pathogens can lead to gastrointestinal illness or other waterborne diseases such as norovirus, giardiasis, and cryptosporidium (Wade et al., 2006). While this issue impacts water bodies in both rural and urban areas, fecal pollution is more prevalent in rural areas due to manure spreading, livestock intrusions into streams, and a higher frequency of septic systems (Nshimyimana et al., 2018; Sekar et al., 2021; Verhougstraete et al., 2015; Wiesner-Friedman et al., 2022; Wilkes et al., 2013). Urban sources of fecal pollution are typically illicit discharges, failing sewage infrastructure, and precipitation events that cause combined or sanitary sewer overflows (Derx et al., 2021; Frick et al., 2020; Hachad et al., 2022; Lyons et al., 2021; Marsalek & Rochfort, 2004).

Water quality standards established by states set limits for fecal pollution, determined by the concentration of fecal indicator bacteria (FIB). *Escherichia coli* (*E. coli)* is the most common FIB in freshwater systems. Traditional methods for measuring FIB are culture-based, which are effective but have several limitations. For example, fecal indicator bacteria are a general measurement of fecal pollution but provide no host-species-specific information, making source identification challenging, especially for non-point source pollution (Harwood et al., 2014). Additionally, *E. coli* are facultative anaerobes and have been known to colonize and reproduce in the sediments and on aquatic vegetation leading to potentially false positive results (Jamieson et al., 2005; Safaie et al., 2021).

Microbial source tracking (MST) can address these limitations by targeting either the host or a host-associated microorganism that is strictly anaerobic and cannot persist or reproduce in the environment. Therefore, their presence indicates recent fecal contamination. This technique uses polymerase chain reaction (PCR) technology to target and amplify DNA sequences indicative of host-specific bacterial groups, typically in the genus *Bacteroides*, that are found in the intestines of particular species of endothermic vertebrates and shed in feces (Reischer et al., 2013). Understanding the source of fecal contamination contributes substantially to making informed decisions on best management practices (BMPs) to reduce levels of contamination (Hinojosa et al., 2020; Shrestha et al., 2020; Steele et al., 2018). This technology has been implemented in modeling techniques for environmental risk assessment such as Quantitative Microbial Risk Assessment (QMRA) and Soil & Water Assessment Tool (SWAT) to assess the concentration at which MST markers indicate a risk to public health (Boehm et al., 2018; Frey et al., 2013). These models have highlighted the importance of associating land-use characteristics, such as soil type, land cover classification, population demographic, and common fecal sources with MST/FIB results, improving source identification. These modeling techniques have been combined with MST results and culture-based monitoring to assess the risks to public health (Sinigalliano et al., 2021).

The United States Environmental Protection Agency (USEPA) developed a standardized method to quantify human fecal contamination using quantitative PCR (qPCR) (US EPA 2019). This PCR-based technique is an improvement over traditional FIB testing, but it is prone to inhibition, has a relatively high lower limit of quantification, and requires the use of a standard curve to obtain concentration estimates (Cao et al., 2015; Doi et al., 2015; Staley et al., 2018). A newer PCR technology called droplet digital PCR (ddPCR) has been developed to reduce these problems.

Droplet digital PCR (ddPCR) is an endpoint reaction that requires no standard curv e and overcomes the majority of limitations of qPCR regarding sample inhibition and sensitivity (Cao et al., 2015). This technique partitions each reaction into thousands of nanoliter-size droplets, each undergoing a separate PCR reaction. These droplets are individually assessed for amplification (via fluorescence), allowing quantification of DNA targets using Poisson statistics and the fraction of positive droplets (Hindson et al., 2011). This new method has been recently employed in several studies to detect species in aquatic environments and perform MST testing (Doi et al., 2015; Pendergraph et al., 2021; Zhu et al., 2020).

The objectives of the present study were to (1) compare the effectiveness of qPCR and ddPCR for MST testing, (2) determine the fecal contaminants impacting two watersheds in western Michigan, and (3) identify watershed characteristics that appear to increase the risk of fecal contamination. The two MST technologies, qPCR and ddPCR, were used to evaluate potential sources of fecal contamination. Fecal pollution was monitored using traditional culture-based quantification of *E. coli*. Three qPCR host-specific genetic markers were employed for MST testing: human (HF183/BacR287), Cow (CowM2), and general ruminant (Rum2Bac) (Green et al., 2014; Mieszkin et al., 2010; Shanks et al., 2008). Quantitative PCR was used in 2019 to measure HF183, and ddPCR was used in 2021 to measure all three markers.

## 2. Methods

### 2.1. Watershed Characterization

Watershed characteristics thought to be associated with FIB sources and their transport were analyzed. Land cover analyses were conducted for both watersheds using the 2019 National Land Cover Database (NLCD; Dewitz et al., 2021). Land cover was aggregated to level 1 Anderson classifications. Hydrologic soil groups and septic field suitability were obtained from the Natural Resources Conservation Service’s (NRCS) SSURGO database (USDA 2022). Population estimates were derived from the 2020 US census block groups for Ottawa County, MI. Depth to water table estimates were provided from Ottawa County (Curtis et al., 2018). Precipitation data was provided by Enviroweather (www.enviroweather.msu.edu). Septic system numbers were estimated using the Microsoft Building Footprints - Tiles dataset (Microsoft 2022). The maps provided by cities and townships, if available, were digitized to derive public sewer service areas. The service area was defined as the political (city or township) boundary if unavailable. These service areas were used as an exclusion mask to eliminate buildings connected to the public sewer system. The remaining buildings were each allocated one septic system, which may overestimate septic prevalence due to the inclusion of unoccupied buildings (e.g., detached garages, barns, etc.). To correct for this, the data set was compared against the Central West Michigan Public Well Records (EGLE 2022). It was assumed that all areas outside of the sewer service area with wells also had a septic system. A key limitation of the well data set was that the entry of wells into this system was voluntary until January 1, 2000. We took the average of both approaches to derive approximate septic system estimates. All watershed characterization was performed in ArcGIS 10.8.1.

### 2.2. Study Sites

This study measured fecal bacteria concentrations at 10 sites on Bass River and 15 sites on Deer Creek selected by the Ottawa Conservation District as representative of each watershed’s tributaries (Figure 2.1). All sites were sampled once per week for five weeks and included two wet-weather events of ≥ 0.25 in (6.35 mm) within 24 hours prior to sampling. If the two wet-weather events did not occur during the initial five-week sample period, up to two additional sampling events were added. Sampling in 2019 occurred in August–September, while the 2021 sampling occurred in June–July.

**Figure 2.1.**
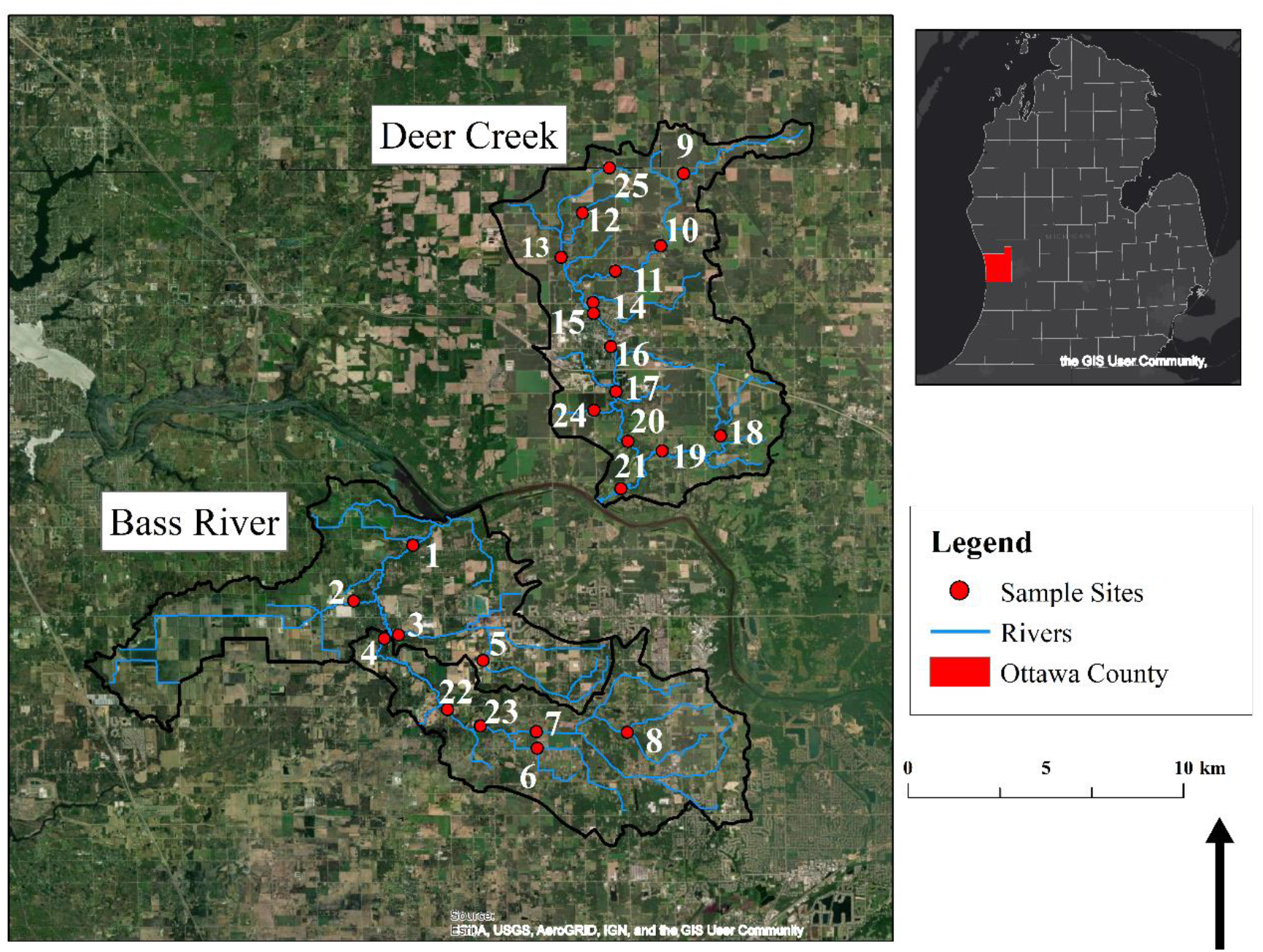
Sample Sites in two watersheds located in Ottawa County Michigan

### 2.3. Sample Collection

Grab samples were collected at three in-stream locations (left, center, right) in the flowing portion of the river, immediately upstream of the road crossing. A 500 mL grab sample was collected at each location by immersing a sterile HDPE bottle below the stream surface, avoiding both surface water and any disturbed sediment. Samples were taken with a collection pole from streams too deep to cross and was decontaminated between sample locations. Samples were kept at 4 °C for transport to the lab for analysis and were processed within 6 hours of initial collection. Two sites were chosen at random for duplicate analysis of all tests to assess lab consistency. Three field blanks consisting of 100 mL of Type 1 DI water were used to ensure no cross contamination occurred in sample transit.

### 2.4. Reference DNA Preparation

Reference DNA materials for qPCR consisted of two plasmids and salmon sperm DNA. The Rose Lab at Michigan State University (MSU) provided the plasmids for internal amplification control (IAC), the sample processing control (SPC), and calibration standards. The calibration standards for qPCR were quantified and linearized by the Rose Lab at concentrations ranging from 10^1^–10^5^ copies/2 μL. The Rose Lab also provided salmon DNA suspended in Qiagen AE buffer at a concentration of 0.02 μg/mL (SAE). The plasmids were stored at −20 °C, and the SAE was stored at 4 °C. Standard Reference Material 2917 (NIST) was used for control sample DNA (Willis et al., 2022). Initial SAE provided by the Rose Lab was diluted to 2 ng/mL for ddPCR. All ddPCR standards were stored at 4 °C.

### 2.5. FIB Quantification and MST Filter Preparation

This study had 789 samples from 2019 and 2021 that were tested for *E. coli*. A subset of 237 samples were analyzed for host specific fecal indicator markers. All samples were quantified for *E. coli* using the Colilert-18® Quanti-Tray/2000® IDEXX system as per manufacturer’s instructions (IDEXX Laboratories, Westport, ME). All samples were initially run at a 10x dilution using phosphate buffered saline (PBS). The Colilert-18® Quanti-Tray/2000® quantification range was >10−24,196 MPN/100 mL. Additional dilutions were used if the sample exceeded the upper quantification limit. All field blanks and two lab blanks were run with each batch as quality control (QC). The geometric mean was taken at each sample site, combining the left, center, and right *E. coli* results, to give a representative *E. coli* concentration. A monthly geometric mean was also calculated for each site.

### 2.6. MST Filtration

100 mL of the center sample was filtered through 0.45 μm PALL MicroFunnel™ Filter Funnels (Pall Corporation, Ann Arbor, Michigan). After filtering, 20 mL of PBS or phosphate buffered water (PBW) was used to rinse the funnel. The filter was collected using aseptic techniques and placed in a 2.0 microfuge tube containing 0.3 g of acid washed glass beads (Sigma). For QC, 20 mL of PBS for qPCR or PBW for ddPCR was passed through a clean filter and collected. Three QC filters were collected for each filtration period. The samples and controls were stored at −80 °C until analysis.

### 2.7. qPCR Methods

DNA extraction for qPCR samples was performed with DNA-EZ RW02 kit spin columns (GeneRite, North Brunswick, NJ) according to manufacturer instructions. Three QC filters were processed with each sample batch as method extraction blanks. Processed samples were stored at 4 °C (<48 hours) before qPCR amplification. Molecular quantification of HF183/BacR287 was conducted using EPA method 1696 (US EPA, 2019) detection method with a Thermo Fisher Scientific StepOnePlus™ (Thermo Fisher Scientific, Grand Island, NY). Analysis was performed on 25-μL reactions using TaqMan™ Environmental Master Mix 2.0. Quantitative PCR reactions contained 23 μL of assay mix which included 12.5 μL of Master Mix, 2.5 μL of 2.0 mg/mL stock solution bovine serum albumin (BSA) from fraction V powder (Sigma B-4287 or equivalent), 1,000 nM primers (forward and reverse), 80 nM of probe, 2.0 μL of IAC plasmid, nuclease free PCR water, and 2.0 μL of the template DNA or DNA calibration standard. HF183/BacR287 was analyzed in duplex with the IAC, and Sketa22 was run as a single plex (Table 3). Sketa22 was used as a SPC in both methods. Each plate was also analyzed with DNA calibration standards (gBlock®) at concentrations ranging from 10^1^–10^5^ target gene copies of reference DNA. Amplification was performed using Thermo Fisher Scientific StepOnePlus™ (Thermo Fisher Scientific, Grand Island, NY) under the following conditions: 50 °C for 2 minutes, 95 °C for 10 minutes, 40 cycles of 95 °C for 15 seconds and 60 °C for 1 minute. The threshold was set at 0.03 for all molecular markers. Quantification values (C_q_) were exported into EPA’s workbook for further analysis (US EPA, 2019). All samples and controls were run in triplicate. Controls included positive controls, extraction blanks, and no-treatment controls (NTCs). These controls were run with each plate to monitor for contamination during sample processing.

**Table 1.**
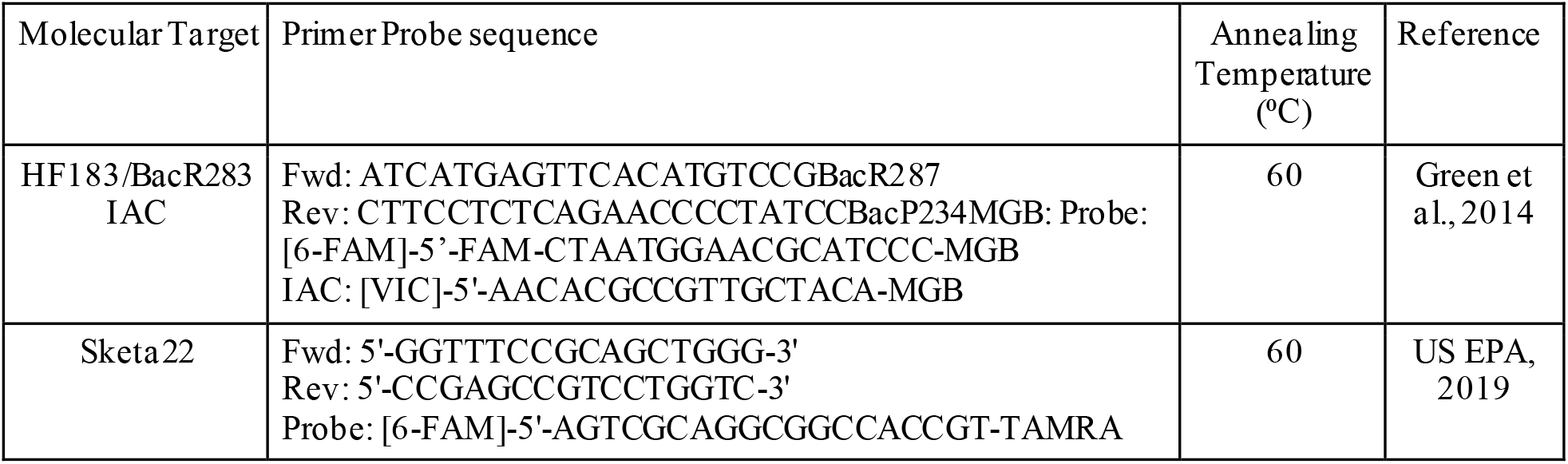
Oligotide sequences for qPCR analysis

**Table 2.**
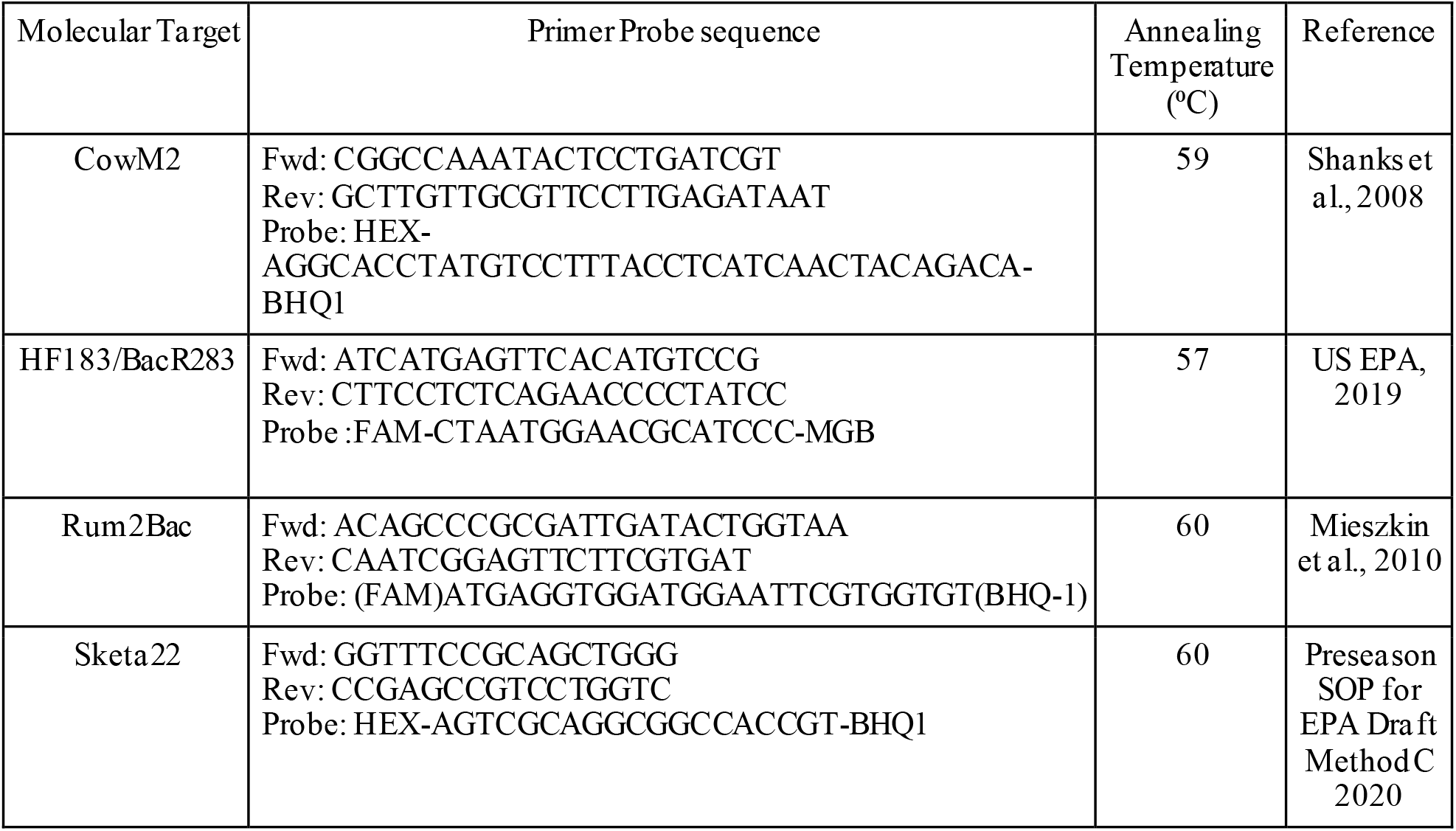
Oligotide sequences for ddPCR analysis

**Table 3.** Land Cover Categories in the Bass River and Deer Creek Watersheds

| Landcover Type | Bass River (%) | Deer Creek (%) |
| --- | --- | --- |
| Open Water | 0.6 | 0.2 |
| Urban | 15 | 13.7 |
| Forest | 16 | 4.5 |
| Agriculture | 53.4 | 72.9 |
| Wetlands | 15 | 8.7 |

Each plate has quality assurance and control (QA/QC) criteria for both qPCR run efficiency and controls to be accepted for analysis. The calibration curve required R^2^ value ≥0.97 and an amplification efficiency of 0.9-1.10. The amplification efficiency, E, is given by equation (2.1).

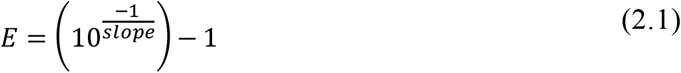

For controls, NTCs and extraction blanks needed to be below the lower limit of quantification (LLOQ), 35.16 C_q_, for all markers except the IAC. All samples spiked with IAC needed to yield results within a C_q_ standard deviation ≤1.16. Finally, the SPC samples required a standard deviation <0.62 C_q_. Samples that passed QA/QC requirements were used to estimate log_10_ copies/reaction using a standard curve with adjustments for DNA recovery provided by the SPC assay results in the Automated Draft EPA Methods 1696 and 1697 Data Analysis Tool in Excel.

### 2.8. ddPCR Methods

DNA extraction for ddPCR was performed using a crude lysis method as described in Aw et. al (2019) with a modification in the initial SAE concentration. Briefly, 600 μL of 0.2 ng/mL salmon was added to each 2 mL microcentrifuge tube and was homogenized at 6.0 m/s for 60 seconds then centrifuged for one minute at 12,000×*g*. Supernatant was taken, placed in a fresh microcentrifuge tube, and clarified through additional centrifugation for 5 minutes. Finally, supernatant was placed in 1.5 mL low-bind microcentrifuge tubes before ddPCR amplification. One extraction blank was processed per batch using a QC filter.

Molecular quantification of host associated microbial source tracking targets was conducted using MSU’s 2021 ddPCR MST method using BIO-RAD QX200™ Droplet Digital PCR (ddPCR) System. 22-μL reactions were made using Bio-Rad’s ddPCR Supermix for Probes (No DUTP) (Bio-Rad Laboratories, Richmond, CA). Each ddPCR reaction contained 16.5 μL of an assay mix with 11 μL of Supermix, 900 nM primers (forward and reverse), 270 nM of probes, nuclease-free PCR water, and 5.5 μL of the template DNA. Targets HF183/BacR283 and Sketa22 were analyzed in duplex, and CowM2 and Rum2Bac were run in duplex (Table 4). Droplets were generated using Bio-Rad AutoDG, and amplification was done using a Bio-Rad C1000 Touch™ Thermal Cycler (Bio-Rad Laboratories) under the following conditions: 95 °C for 10 minutes and 40 Cycles of (94 °C for 30 seconds, 58 °C for 1 minute). All ddPCR reactions and controls were analyzed in triplicate. Each ddPCR plate contained positive, negative, NTCs for each assay mix. Analysis was performed on Bio-Rad QX200. All samples were spiked with salmon DNA as a sample processing control (SPC) to assess sample inhibition.

**Table 4.** Hydrologic Soil Groups in the Bass River and Deer Creek Watersheds

| Hydrologic Soil Group | Bass River (%) | Deer Creek (%) |
| --- | --- | --- |
| A | 34 | 6 |
| A/D | 35 | 11 |
| B | 1 | 5 |
| B/D | 6 | 6 |
| C | 2 | 18 |
| C/D | 11 | 20 |
| D | 9 | 30 |

QA/QC were performed for all samples. Samples with less than 10,000 accepted droplets were excluded from data analysis. Thresholds were manually set for all wells simultaneously with the criteria of being above the negative control and NTC wells, and below the positive control wells. All samples were spiked with salmon DNA to test for inhibition. Detection of at least three positive droplets in one of the triplicate wells provided evidence that the DNA target was present in the sample. The minimum detection limit (MDL) was 360 gene copies/100 mL. Results were calculated as gene copies per 100 mL (GC/100 mL) using equation 2.2.

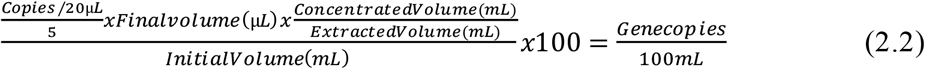

### 2.9. Statistical Analysis

The R programming language and computing environment, version 4.1.3 (R Core Team, 2021), was used for statistical analysis. The Colilert geometric means for the two years at each sampling site in the Bass River and Deer Creek were treated as matched pairs and a nonparametric Wilcoxon signed-rank test was performed (Hollander et al. 2014). The null hypothesis of no difference was tested against the two-sided alternative hypothesis at a significance level of α = 0.05. The effectiveness of qPCR and ddPCR was assessed and compared by calculating the maximum-likelihood estimate of the detection probability and its 95% confidence interval (CI) using Wilson’s confidence interval (Brown et al., 2001). All watershed characteristics and FIB data were assessed for pairwise relationships between variables followed by non-parametric quantile regression. Visual inspection of the quantile regression suggested linear relationships between only CowM2 and hydrologic soil groups A and D, but the slopes were not significantly different from zero (p > 0.05).

## 3. Results

### 3.1. Watershed Characteristics

#### 3.1.1. Bass River

Bass River (HUC: 04050006-0706; -0707) is a rural 12,958 ha watershed with an average slope of 5% located in Ottawa County, MI. This watershed has a population of ∼23,000 and ∼62% are on septic systems (∼2,500 septic systems). Land use is primarily agricultural (53.4%) with forest and wetland areas fragmented throughout the watershed (16% and 15%, respectively) (Table 3, Figure 3.1). Urban land use is primarily low intensity and occurs mainly along roads (15%) (Table 3, Figure 3.1).

**Figure 3.1.**
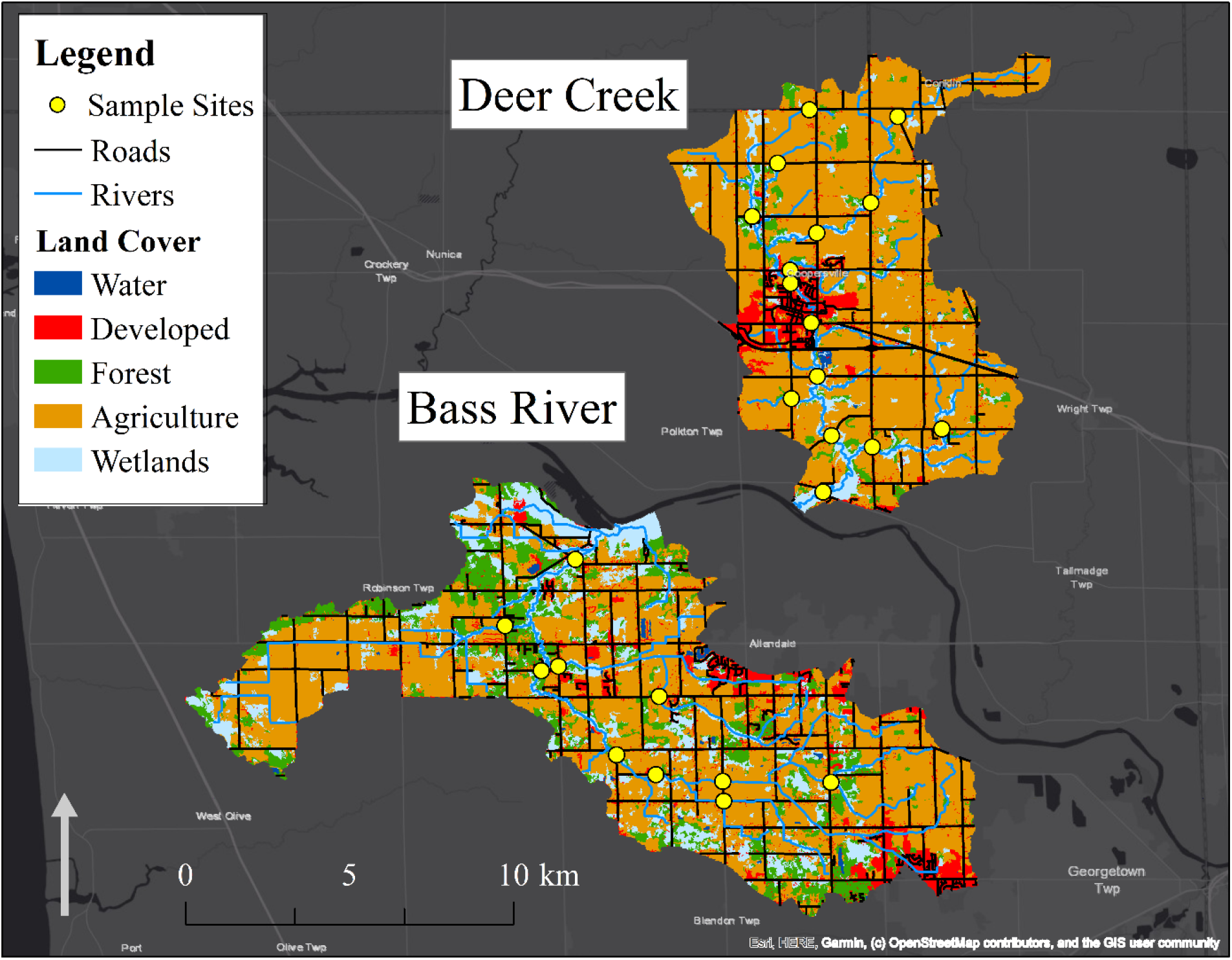
Bass River and Deer Creek Land Cover characterization

Bass River watershed contains sandy, low to moderately textured soil with high infiltration rates and low runoff potential. Most of the watershed falls into the A-B hydrologic soil group (70%) (Table 4, Figure 3.2). While these soils normally indicate good infiltration, approximately 40% of soils fall in the A/D-B/D soil type, indicating poor infiltration in undrained areas. Over 97% of the watershed is classified as very limited for septic field suitability in the SSURGO database, indicating high potential for septic failure. Due to extensive agricultural usage, a high-water table, and low septic rating, it is hypothesized that the primary sources of fecal pollution stem from farm practices such as manure spreading during the spring and fall, high utilization of tile drainage, livestock density (cattle), and failing septic systems.

**Figure 3.2.**
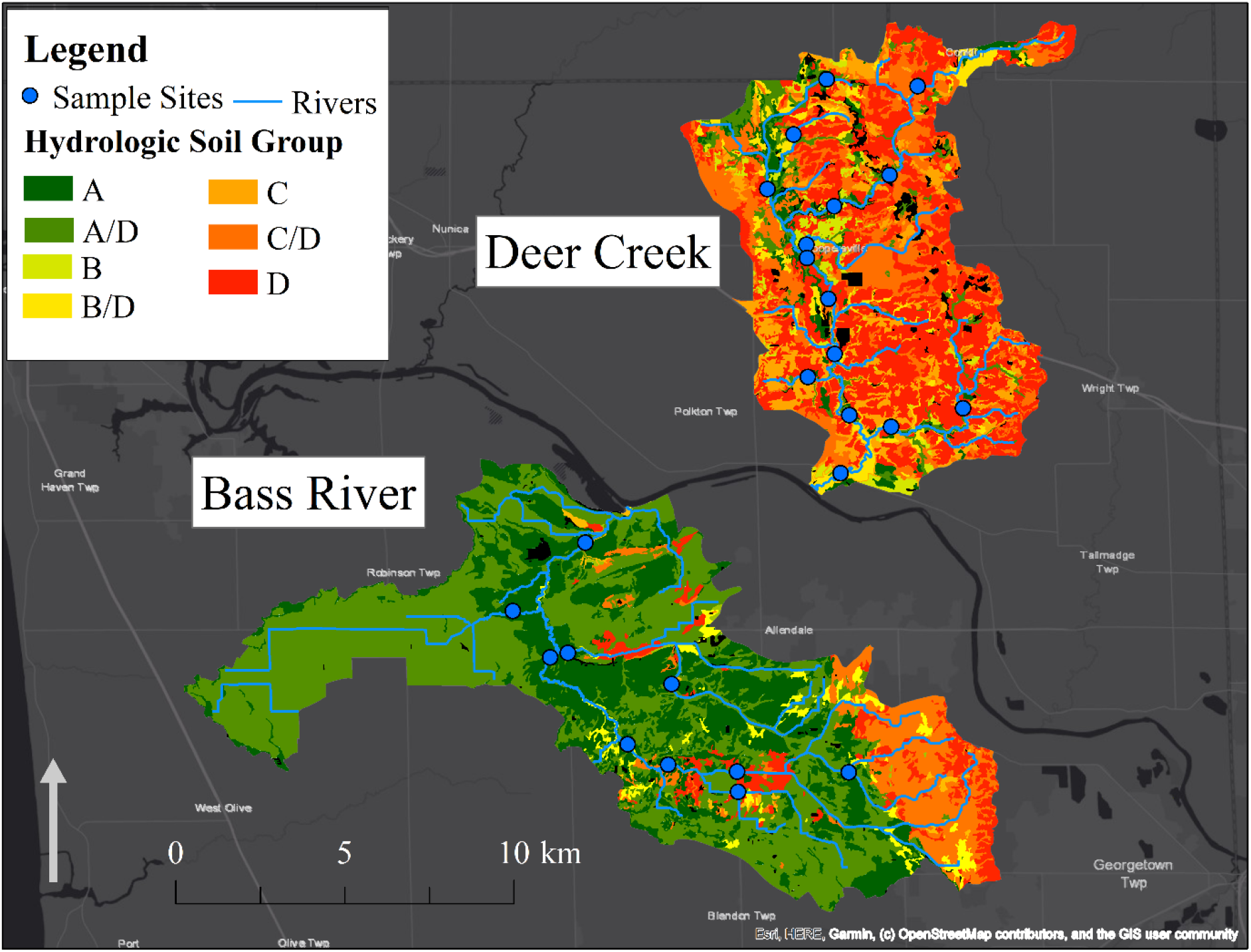
Bass River and Deer Creek Hydrologic Soil Groups

#### 3.1.2. Deer Creek

Deer Creek Deer Creek (HUC: 04050006-0704) is a predominantly rural 9,050-hectare watershed with an average slope of 4% located in the Grand River watershed in Ottawa and Muskegon Counties. This watershed has a population of ∼9,000 people with ∼42% on septic systems (∼1,000 septic systems). Land use is primarily agricultural (72.9%) with urban land use (13.4%) mainly located in the City of Coopersville (Table 3, Figure 3.1). Forest areas are located sporadically throughout the watershed (4.5%), and wetlands are located primarily on the main channel of the watershed and the headwaters (8.7%) (Table 3, Figure 3.1). Deer Creek soil consists of fine clay, providing limited infiltration and high runoff potential. Most of the watershed is in the C-D hydrologic soil groups (68%) (Table 4, Figure 3.2). According to the SSRGO, over 95% of this watershed is classified as very limited suitability for septic fields, indicating a high probability of septic failure. Due to similar agricultural influences as Bass River and low infiltration soil types, it is hypothesized that the main source of fecal pollution stems from farming practices.

#### 3.1.3. Precipitation

The total precipitation during the 2019 sample collection period was ∼186 mm (storm event average 9.3 mm), and the total precipitation during the 2021 sample collection period was ∼202 mm (storm event average 11.2 mm) (Enviroweather).

### 3.2. *E. coli* Results

The mean *E. coli* concentration in Bass River was 1,207 MPN/100 mL in 2019 and 1,734 MPN/100 mL in 2021 (Figure 3.3). The mean *E. coli* concentration in Deer Creek was 1,784 MPN/100 mL in 2019 and 1346 MPN/100 mL in 2021 (Figure 3.3). The grand monthly geometric mean of all sites indicated that the majority of sites exceeded the state partial body recreation standard (1,000 MPN/100 mL) (Part 4 rules, R 323.1062 Microorganisms, promulgated under Part 31, Water Resources Protection, of the Natural Resources and Environmental Protection Act, 1994 PA 451, as amended). A Wilcoxon signed rank exact test showed no statistically significant between-year difference in *E. coli* concentrations in Bass River (p = 0.28, V = 16) or Deer Creek (p = 0.27, V = 62).

**Figure 3.3.**
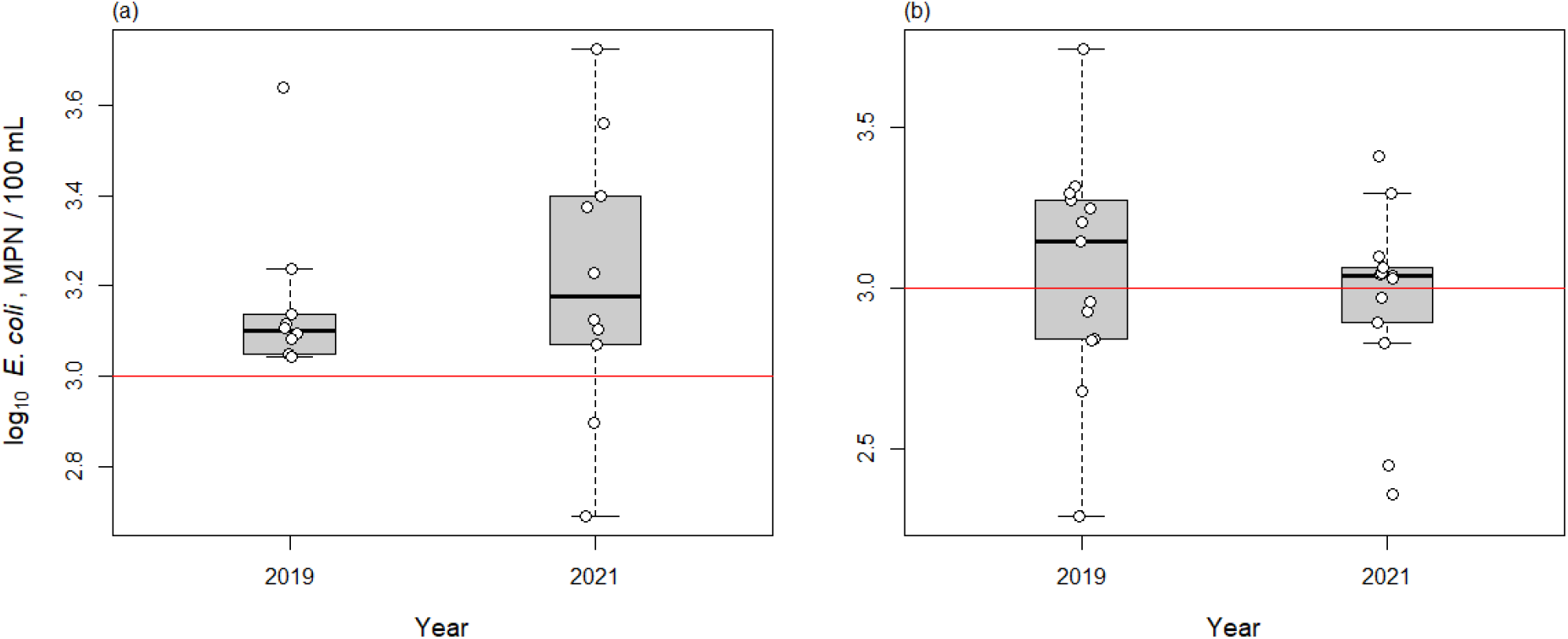
Log_10_ transformed average E.coli concentration in Bass River (a) and Deer Creek (b). The red line represents the partial body recreation standard = 1,000MPN/100 mL

### 3.3. MST Results

All samples passed QA/QC on both methods and indicated no inhibition present. In 2019, 109 samples were tested for HF183/BacR287 using qPCR. HF183/BacR287 was detected in 27% (29 of 109) of these samples, and only 7% of the samples (2 in Bass River and 6 in Deer Creek) were within the quantifiable range (Figure 3.4, Table 5, Table 6).

**Figure 3.4.**
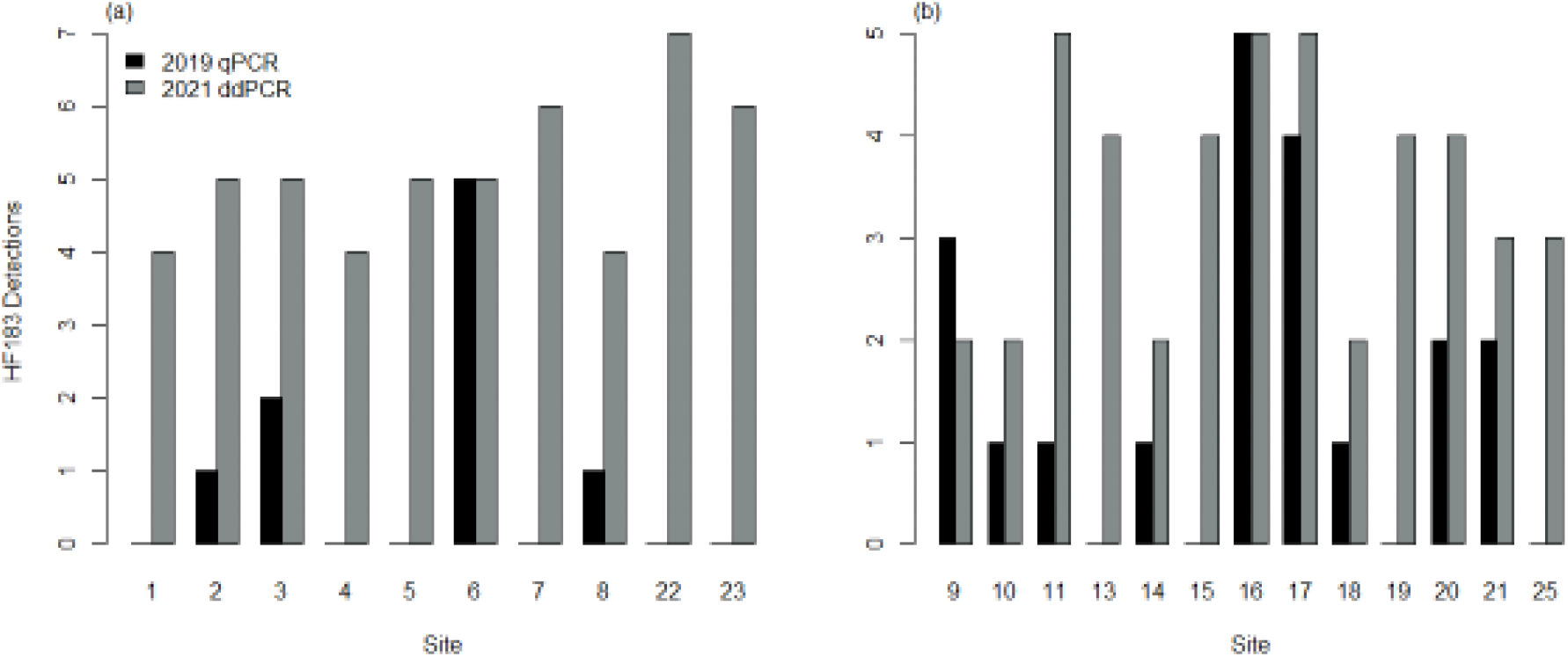
HF183/BacR287 Detection rate for qPCR vs. ddPCR in Bass River (a) and Deer Creek (b)

**Table 5.**
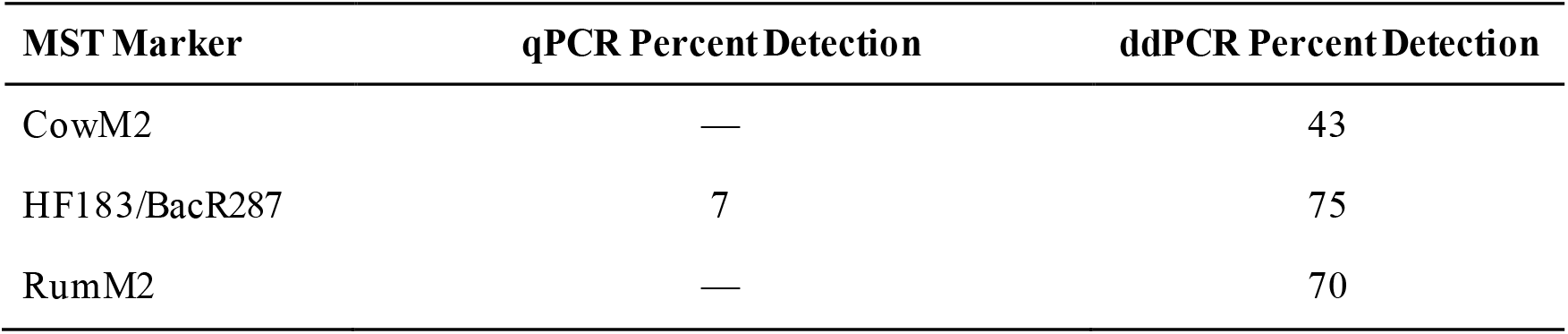
Detection Rates of quantifiable MST Markers in 2019 (109 samples) and 2021 (128 samples)

**Table 6.** qPCR vs. ddPCR detection rate

| Year Sampled (n) | Success Criterion | Percent Detection (95% CI) |
| --- | --- | --- |
| 2019 (109) | qPCR above LLOQ | 7 (4-14) |
| 2019 (109) | All qPCR Detections | 27 (19-36) |
| 2021 (128) | ddPCR above MDL | 75 (67-82) |

In the 2021 season, 124 samples were tested for three MST targets using ddPCR. 95% of the samples had a detection for one or more of the markers; CowM2, HF183/BacR287, and Rum2Bac were detected in 43%, 75%, and 70% of samples, respectively (Table 5). In Bass River, the sites with the highest average signal for CowM2, HF183, and Rum2Bac were sites 22, 6, and 23, respectively (Figure 3.5, Table 7). In Deer Creek, the sites with the highest average signal for CowM2, HF183, and Rum2Bac were sites 17, 16, and 10, respectively (Figure 3.5, Table 8). When comparing the two years of sampling, 57% (13 of 23) of sites tested positive for HF183 in both 2019 and 2021: 4 sites in Bass River and 9 sites in Deer Creek (Figure 3.4).

**Figure 3.5.**
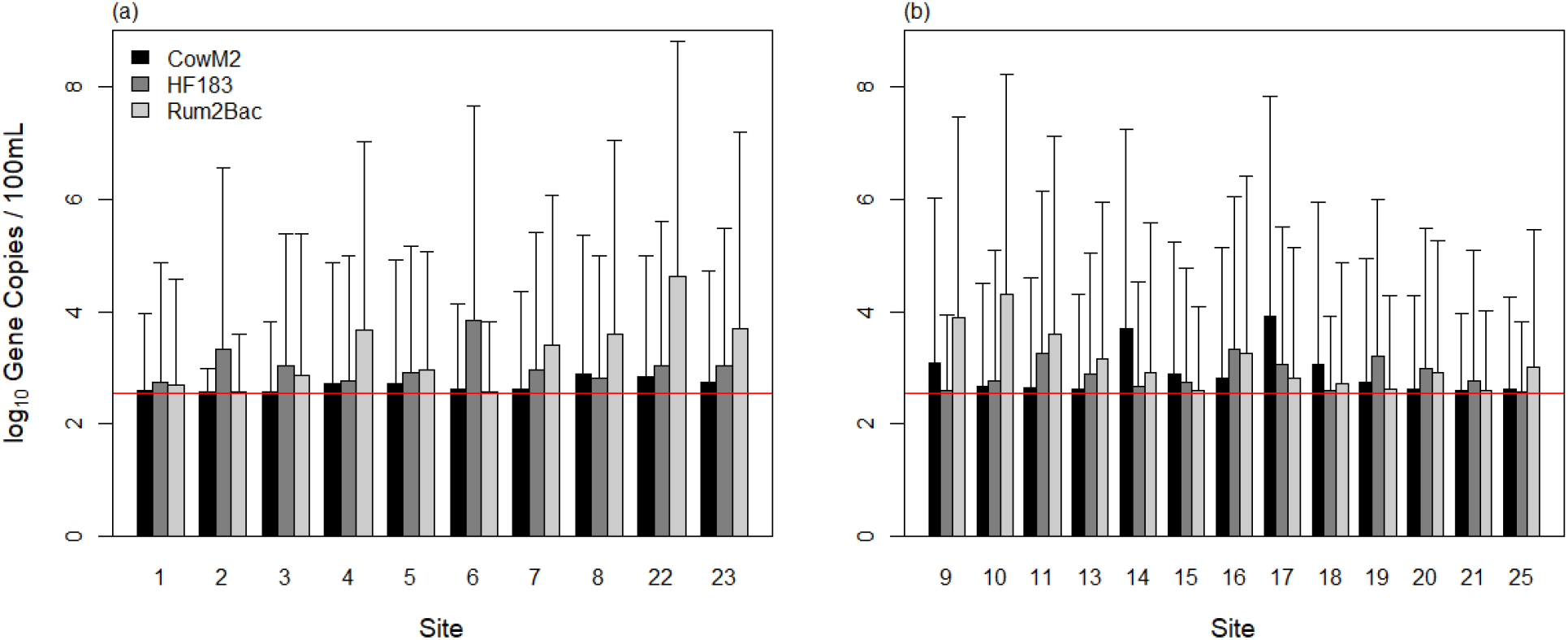
Log_10_ transformed Average MST marker detection in Bass River (a) & Deer Creek (b) by marker. The red line represents the minimum detection limit (MDL) = 360 GC/100ml

**Table 7.** Bass River average molecular markers signal in GC/100 mL

| Site | CowM2 | HF183/BacR287 | Rum2Bac |
| --- | --- | --- | --- |
| 1 | 393 | 563 | 496 |
| 2 | 363 | 2,145 | 371 |
| 3 | 377 | 1,059 | 716 |
| 4 | 516 | 585 | 4,652 |
| 5 | 520 | 819 | 907 |
| 6 | 419 | 6,995 | 379 |
| 7 | 415 | 903 | 2,581 |
| 8 | 780 | 641 | 4,020 |
| 22 | 694 | 1,063 | 41,391 |
| 23 | 544 | 1,088 | 4,911 |

**Table 8.** Deer Creek Average Molecular marker signal in GC/100 mL

| Site | CowM2 | HF183/BacR287 | Rum2Bac |
| --- | --- | --- | --- |
| 9 | 1,229 | 384 | 7,571 |
| 10 | 472 | 579 | 20,206 |
| 11 | 448 | 1,845 | 3,986 |
| 13 | 426 | 752 | 1,437 |
| 14 | 5,078 | 460 | 838 |
| 15 | 779 | 544 | 394 |
| 16 | 634 | 2,082 | 1,813 |
| 17 | 8,443 | 1,170 | 645 |
| 18 | 1,160 | 390 | 523 |
| 19 | 542 | 1,585 | 408 |
| 20 | 425 | 964 | 836 |
| 21 | 384 | 580 | 386 |
| 25 | 405 | 378 | 1,030 |

## 4. Discussion

### 4.1. Land Use and MST characterization

When combined with land use and traditional culture-based monitoring, microbial source tracking provided additional information on the source of pathogen pollution. Verhougstraete et al (2015) analyzed land use, *E. coli*, hydrologic flow, and a human marker, *Bacteroides thetaiotaomicron* (*B. theta*), in a CART (classification and regression trees) analysis to assess human fecal bacteria concentrations in Michigan streams. Their analysis indicated that the number of septic systems significantly correlated with *B. theta* concentrations. They determined that watersheds with over 1,621 septic systems exhibited significantly higher *B. theta* (Verhougstraete et al., 2015), but there was no correlation between system density and *B. theta*. Although a different human MST marker was applied in the present project, the results were consistent with the conclusions drawn by Verhougstraete et. al. (2015). Sowah et al. (2017) compared areas with a low density of septic systems (< 38 per km^2^) to areas with a high density of septic systems (> 77 septic systems per km^2^) to assess the impact on groundwater recharge through both baseflow and wet weather conditions. They concluded that areas with >87 septic systems per km^2^ had negative impacts through groundwater contributions and failing units (Sowah et al., 2017). Bass River contained a higher number of septic systems (> 1,621) and higher average concentrations of the human marker when compared to Deer Creek, which had fewer septic systems. The increased number/density of septic systems was found to be associated with higher detection of human fecal markers in a variety of studies in both temperate and tropical climates (Dila et al., 2018; Jent et al., 2013; Wiesner-Friedman et al., 2022).

Studies of urban watersheds found that precipitation caused both increased flow and an initial increase in human fecal markers just before the peak hydrograph measurement, with a subsequent decrease due to dilution, leading to a lower average signal (McLellan et al., 2018). This was seen in both watersheds with separate sanitary and stormwater sewage as well as areas with combined sewer overflows. These studies concluded that continued detection of human pollution during baseflow indicated illicit discharge, both intentionally to stormwater lines, and unintentionally through failing infrastructure. While the majority of our sites are outside of sewer service, several sites in Deer Creek are located in the service area and displayed elevated levels of the human fecal marker during baseflow and high-flow events, suggesting illicit discharges or failing infrastructure.

In the rural areas, agriculture is a primary source of fecal contamination. Previous studies have shown a positive association between the detection of livestock markers, elevated runoff, and livestock intrusion (Ballesté et al., 2020; Dila et al., 2018; Jent et al., 2013; Wilkes et al., 2013). Another study sampled a variety of watersheds located in the Great Lakes region and found that cow and general ruminant markers had a positive correlation with cattle density, increased presence during runoff events, and areas with a high proportion agricultural tile drainage (Dila et al., 2018). Additional studies focusing on livestock intrusion, tile drainage in agricultural fields, and runoff from agriculturally dominant watersheds support these conclusions (Ballesté et al., 2021; S. K. Frey et al., 2015; Wilkes et al., 2013). The present study is consistent with these findings; both watersheds displayed a high prevalence of ruminant pollution, most likely related to high livestock densities and widespread use of tile drainage. In addition to basic land use classification, a variety of other studies in both tropical and temperate environments worldwide, have made associations between microbial indicators, hydrology, water chemistry, and land use characteristics to identify sources of fecal pollution in a variety of water sources, streams, groundwater, and river systems, (Frick et al., 2020; Jent et al., 2013; Lyons et al., 2021; Sekar et al., 2021; Steinbacher et al., 2021; Wiesner-Friedman et al., 2022). These studies highlight the importance of connecting pathogen contamination with other land use and chemical characteristics to confirm the identification of non-point sources of fecal contamination.

In this study, we applied land use characteristics, FIB, and MST results of two watersheds Bass River and Deer Creek in the Grand River watershed, to assess potential sources and sites of fecal pollution. Based on watershed characteristics, we determined that human, cow, and general ruminant were the most likely sources of fecal pollution. Our *E. coli* results and precipitation records over the two sampling periods indicated no significant difference between sample period or location. Over half of the sample sites had repeat detections of the human marker. While the *E. coli* results showed no significant between-year difference in pathogen concentration, MST results indicated an increase in pathogen detection from 2019-2021 (27%-75%). This increase could be due to improved PCR technology through ddPCR quantification.

### 4.2. PCR Methodology Comparison

The *E. coli* concentrations between the two seasons were not statistically different, and it was therefore expected that the fecal marker distribution would be similar for both years. However, when measuring the human marker, HF183, the 2019 results from qPCR indicated that 73% of samples were below the MDL, indicating the absence of the human marker. When measuring the same marker with ddPCR in 2021, only 25% of samples were below the MDL (Table 6). One explanation for this discrepancy could be due to differences in precipitation during sampling periods. Previous studies have indicated a link between increased precipitation resulting in increased leachate from failing septic systems (Dila et al., 2018; Frick et al., 2020; Jent et al., 2013; Verhougstraete et al., 2015). While there was no evidence indicating a between-year difference in *E. coli* concentrations or precipitation totals (∼19 cm in 2019, ∼20 cm in 2021), minor differences in precipitation may account for the variation in presence of the human marker between sampling years. Another possible cause for the difference in detection is the sensitivity of the technology used (qPCR vs. ddPCR). This limits our study since samples were not analyzed on both technologies each year to make a direct comparison.

Despite these limitations, our results comparing PCR technologies agree with previous studies that analyzed split samples with both methods (Cao et al., 2015; Yang et al., 2014). Additional studies indicated that ddPCR had higher sensitivity than qPCR and provided absolute quantification of the DNA target, which increased the reproducibility of quantitative results (Hindson et al., 2011; Zhao et al., 2016). While these factors highlight the advantages of ddPCR compared to qPCR, our study demonstrates the usefulness of qPCR as an analytical tool. qPCR is effective in areas with high amounts of bacterial pollution because it has a greater upper limit of quantification than ddPCR. However, the higher sensitivity of ddPCR is needed in areas where the markers of interest are in low concentrations. Additional factors such as the cost to implement, number of markers, and amount of pollution expected also may impact the choice of PCR platform (Nshimyimana et al., 2019).

### 4.3. Implications for the watersheds

#### 4.3.1. Bass River

The primary sources of pollution in Bass River were ruminants and humans. The human marker was detected at all sites, 40% had repeat detections, and 50% had concentrations over 1,000 GC/100mL (Figure 3.3; Table 7). When looking at the 2021 results, small amounts of the cow-specific marker were detected during the sample season, indicating that they are not the main source of fecal contamination in Bass River (Table 7). Site 6 showed exceptionally high amounts of HF183 at 6,995 GC/100 mL. This is likely due to the presence of multiple septic systems and it is located in a type A/D hydrologic soil group surrounded by type D soils (Figure 3.2). These watershed characteristics indicate that this site may have poor infiltration, increasing the leaching from failing septic systems into the stream. Future remediation efforts should focus on locating and replacing failing septic systems that contribute to this pollution and any potential illicit discharges that may be present in areas with sewage hookups. The ruminant marker was detected at all sites with 50% of sites having over 1,000 GC/100 mL of (Table 7). Sites 4, 22, and 23 showed elevated concentrations of the ruminant marker, with concentrations over 4,000 GC/100 mL. All three of these sites are located at the edge of forested sites next to agriculture (Figure 3.1). Due to these watershed characteristics, the source of ruminant pollution could be an animal other than cows, such as whitetail deer or other free-range ruminant livestock such as goats or sheep. This information indicates that future remediation efforts should focus on BMPs that target human and ruminant sources of fecal pollution. Future research in this watershed should continue monitoring for human markers but also include other potential ruminant hosts of interest such as whitetail deer, sheep, goats.

#### 4.3.2. Deer Creek

Our results from both sampling seasons suggest that the majority of fecal pollution stemmed from human and ruminant sources. The human marker was detected in 69% of samples from both sample years, and 30% of the sites had a concentration over 1,000 GC/100 mL (Figure 3.3; Table 8). The cow marker was detected in higher concentrations in Deer Creek than in Bass River, with over 30% of sites exceeding 1,000 GC/100 mL (Table 8). Sites 14 and 17 had the highest concentrations of CowM2, with over 5,000 GC/100 mL. This is most likely due to land use, as site 14 is located between urban and agricultural landcover (Figure 3.1), and site 17 in an agriculturally dominant area. Both sites have relatively low infiltration (A/D-B/D at site 14 and C and D at site 17), which increases the runoff potential (Figure 3.2). Due to their locations, both may also be receiving runoff from nearby tile-drained agricultural fields leading to elevated cow concentrations. Site 16 had the highest concentration of HF183 in Deer Creek over the sample period, with a concentration of 2,082 GC/100 mL. This sample site is located in urban land cover and in an area of Deer Creek served by municipal sewer. Based on the land use characteristics and low infiltration hydrologic soil groups surrounding the site, this elevated signal is probably due to illicit discharges or failing infrastructure (Figure 3.1, 3.2). The general ruminant marker had the highest average concentration, with 46% of sites at a concentration over 1,000 GC/100 mL (Table 8), and sites 9,10, and 11 had concentrations over 3,500 GC/100 mL. All three sites are surrounded by agricultural land use, have non-porous hydrologic soil groups, and are located on the same tributary (Figure 3.1, 3.2). Because of this, it can be assumed that most ruminant pollution originates from field runoff and livestock intrusion. This elevated marker signal could also originate from other ruminants such as whitetail deer, sheep, or goats. Further testing should focus on additional host-specific MST ruminant targets such as deer, sheep, and goats. Future remediation efforts should focus on BMPs that keep manure and livestock out of streams.

## 5. Conclusions

Microbial source tracking in conjunction with FIB monitoring and watershed characteristics improved the identification of non-point source fecal contamination in two watersheds in Ottawa County, MI USA. Our study indicated ddPCR had higher sensitivity for the detection of DNA targets in environmental samples compared to qPCR, making it a better tool for the detection of multiple targets or highly diluted samples. This study highlights some key considerations when choosing a PCR platform to best suit stakeholder goals. The number of septic systems and soil type had an association with the detection of MST targets from both anthropogenic and non-anthropogenic sources in rural watersheds. When land use and soil data are combined with MST, stakeholders are provided a more insightful assessment of nonpoint sources. This additional information allows them to make more informed decisions regarding best management practices and revision of total maximum daily loading limits. This procedure is applicable to a variety of waterbodies (rivers, streams, and lakes), in a variety of climates worldwide.

## Acknowledgments

We thank Jacob Weston for providing the initial septic estimation data, Yingqing Deng and Noah Cleghorn for their assistance with the collection and analysis of the 2019 samples, an d Alexis Porter, Katelyn Anderson, and Brian Scull for reviewing article drafts. We also would like to thank the Ottawa Conservation District, EGLE/EPA 319 Grant 2018 -0021; Grand Valley State Universities Presidential Research Grant; and The West Michigan Air & Waste Management Association’s individual scholarship for funding this project. This work was done in partial fulfillment of requirements for a Master of Science in Biology at the Robert B. Annis Water Resources Institute.

## Author Contributions

RR and BJ designed the study and secured funding. JJH, and RR performed the study. JJH, MNJ, JNM, SAW, and RR analyzed the results of the study. JJH wrote the article. MNJ, JNM, SAW, BJ and RR commented on draft versions of the article. All authors have approved the final article.

## Notes

### Competing Interest Statement

The authors have declared no competing interest.

### Summary of Updates

Added Conclusion section

